# Evaluation of three sets of advanced backcrosses of eggplant with wild relatives from different genepools under low N fertilization conditions

**DOI:** 10.1101/2023.05.04.539369

**Authors:** Gloria Villanueva, Mariola Plazas, Pietro Gramazio, Reyes D. Moya, Jaime Prohens, Santiago Vilanova

## Abstract

The development of new cultivars with improved nitrogen use efficiency (NUE) is key for implementing sustainable agriculture practices. Crop wild relatives (CWRs) provide valuable genetic resources for breeding programs aimed at achieving this goal. In this study, three eggplant (*Solanum melongena*) accessions together with their advanced backcrosses (ABs; BC3 to BC5 generations) with introgressions from the wild relatives *S. insanum*, *S. dasyphyllum*, and *S. elaeagnifolium* were evaluated for 22 morpho-agronomic, physiological, and N use efficiency (NUE) traits under low nitrogen fertilization conditions. The three *S. melongena* recurrent parents were also evaluated under control (normal) N fertilization. Reduction of N fertilization in the parents resulted in decreased chlorophyll content-related traits, aerial biomass, stem diameter and yield, and increased NUE, nitrogen uptake efficiency (NUpE), and nitrogen utilization efficiency (NUtE). However, the decrease in yield was moderate, ranging between 62.6% and 72.6%. A high phenotypic variation was observed within each of the three sets of ABs under low nitrogen conditions, with some individuals displaying improved transgressive characteristics over the recurrent parents. Genotyping with the SPET 5k probes platform revealed a high, although variable, degree of recovery of the recurrent parent genome in the ABs and allowed the identification of 16 QTLs. Different allelic effects were observed for the introgressed QTL alleles. Several candidate genes were identified in the QTL regions associated with plant growth, yield, fruit size, and NUE-related parameters. Our results show that eggplant materials with introgressions from CWRs can result in a dramatic impact in eggplant breeding for a more sustainable agriculture.

## 1. Introduction

Enhancing crop productivity is a fundamental objective in agriculture and remarkable advancements have been achieved in this area since the beginning of the 20^th^ century. This has been accomplished, in part, through the widespread utilization of nitrogen (N) as a fertilizer^1^. However, excessive use of N fertilization can lead to negative environmental impacts, such as groundwater and surface water contamination, loss of biodiversity, increased greenhouse gas emissions, and ozone layer depletion^2–4^. In addition, synthetic N fertilizers require large amounts of energy to be produced^5^. Therefore, to mitigate these negative consequences, selection and development of new varieties with improved crop nitrogen use efficiency (NUE) is a major objective of plant breeding for a more sustainable agriculture^6, 7^.

The utilization of plant genetic resources is essential for implementing breeding programs to address challenges associated with changes in climatic conditions. In this context, crop wild relatives (CWRs) are of great relevance, as they possess inherent adaptations to a wide range of adverse natural conditions^8^. However, the direct utilization of CWRs in breeding programs is often impractical due to the presence of unfavorable traits and genetic barriers. Therefore, the development of advanced backcrosses (ABs) is a viable breeding strategy that expands the available genetic diversity by incorporating genomic fragments of CWR genomes into a mostly cultivated genetic background^9^. Furthermore, ABs are useful for the detection of Quantitative Trait Loci (QTLs) by associating phenotypic variation with specific regions of the genome.

Eggplant (*Solanum melongena* L.), also known as aubergine or brinjal, is a widely cultivated vegetable crop, belonging to the subgenus *Leptostemonum* of the Solanaceae family^10^. It is one of the most important solanaceous crops, ranking second only to tomato (*S. lycopersicum* L.)^11^. The recent development of genomic tools specific for eggplant, such as high-throughput genotyping platforms^12^ and high-quality eggplant genome assemblies^13–16^, among others, has facilitated genomic studies on this crop.

Eggplant wild relatives (CWRs) are classified into the primary (GP1), secondary (GP2) and tertiary (GP3) genepools, based on their level of crossability with the cultivated species. Interspecific hybrids, advanced backcrosses (ABs) and introgression lines (ILs) have been obtained by utilizing a several of these CWRs^17–19^.

Among the CWRs for which ABs have been developed, *Solanum insanum* L. belongs to GP1 and is considered the wild ancestor of the common eggplant (*S. melongena*). This species grows in a wide range of environmental conditions, including infertile soils, and is naturally distributed throughout south and southeast Asia, Madagascar and Mauritius^20^. Among the many eggplant secondary genepool (GP2) species, *S. dasyphyllum* Schumach. & Thonn is part of the Anguivi clade of the *Leptostemonum* subgenus and is considered the wild progenitor of the gboma eggplant (*S. macrocarpon* L.), an African cultivated eggplant^21^. Some studies have shown that *S. insanum*, *S. dasyphyllum* and their interspecific hybrids with eggplant exhibit enhanced drought^22, 23^ and salinity tolerance^24–26^. Another CWR of interest is the American species *S. elaeagnifolium* Cav., which is native to Northern Mexico and the United States and that can thrive in a wide range of climatic conditions, including semiarid areas, being a globally invasive plant^27, 28^. The development of backcrosses of *S. elaeagnifolium* with eggplant has been reported, making available a previously unexploited genepool for eggplant breeding^29^. Additionally, *S. elaeagnifolium* is a potential source for developing new varieties with enhanced drought tolerance ^30^ and adaptation to low N-inputs^31^.

In the present work, we evaluated morpho-agronomic and composition traits of three *S. melongena* accessions (MEL5, MEL1 and MEL3) under two N fertigation conditions and three sets of advanced backcrosses (ABs) of these three accessions with introgressions from eggplant wild relatives *S. insanum*, *S. dasyphyllum* and *S. elaeagnifolium* under low N conditions. The study results provide valuable information in the identification of potential materials for eggplant breeding under low N fertilization. Furthermore, detection of QTLs was made possible through the association of phenotyping data and the availability of high-density genotyping data of the ABs individuals.

## 2. Materials and Methods

### 2.1. Plant material

Three *S. melongena* accessions (MEL5, MEL1 and MEL3) and three sets of advanced backcrosses (AB) of these accessions with, respectively, the eggplant wild relatives *S. dasyphyllum* DAS1, *S. elaeagnifolium* ELE2 and *S. insanum* INS1 were used for the current study^19^. For the set of *S. insanum* ABs (INS1 x MEL5), 25 AB individuals were used, of which eight were from the BC5 and 17 from the BC4S1 generations. In the case of the *S. dasyphyllum* ABs set (MEL1 x DAS1), a total of 59 individuals were used, 41 of them being from the fifth backcross generation (BC5) and 18 of the first selfing of the fourth backcross generation (BC4S1). Finally, for the *S. elaeagnifolium* ABs set (MEL3 x ELE2), 59 individual ABs were also used, of which 16 were from the third backcross generation (BC3) and 43 were from the fourth backcross generation (BC4).

### 2.2. DNA extraction and genotyping

Extraction of genomic DNA of the three recurrent parents and advanced backcrosses individuals was performed following the SILEX DNA extraction method^32^. Isolated DNA was evaluated for quality and integrity by 0.8% agarose gel electrophoresis and spectrophotometric ratios 260:280 and 260:230 and quantified by a Qubit^®^ 2.0 Fluorometer (Thermo Fisher Scientific, Waltham, MA, USA). Diluted DNA samples were genotyped using the eggplant 5k Single Primer Enrichment Technology (SPET) platform consisting of 5,093 probes^12^. Single nucleotide polymorphisms (SNPs) were filtered with Tassel software (version 5.2 Standalone; ^33^) by using a minimum count value of 97%, a minimum allele frequency (MAF) higher than 5%, a maximum heterozygosity proportion of 70% and a minimum distance between adjacent sites of 2,000 pb. After filtering, the number of discriminant SNPs between parents was 826, 1,195, 2,114 and for *S. insanum*, *S. dasyphyllum*, and *S. elaeagnifolium* ABs, respectively.

### 2.3. Cultivation conditions

Plants were grown during the summer season (July to October 2020) in an open field plot located on the campus of the Universitat Politècnica de València (GPS coordinates: latitude, 39° 28’ 55” N; longitude, 0° 20’ 11” W; 7 m a.s.l.). Advanced backcrosses individuals and recurrent parentals lines were randomly distributed in 17 L pots with coconut fiber, spaced 150 cm between rows and 70 cm within rows. Irrigation and fertilization were applied with a drip irrigation system.

The recurrent parents *S. melongena* MEL5, MEL1 and MEL3 were cultivated under two different nitrogen fertilization conditions, namely low (LN) and normal nitrogen (NN) treatments. Seven plants of each *S. melongena* accession together with ABs individuals of each set were cultivated under LN conditions, while seven plants of each *S. melongena* were cultivated under NN conditions.

A physicochemical and composition analysis of coconut fiber was performed before the transplant. Parameters were evaluated following the procedures described in van Reeuwijk^34^ and are shown in Table S1. A chemical composition analysis of water was performed before adding fertilizers. The intake water was slightly basic with low content of nitrates, nitrites, phosphates, ammonium, magnesium and potassium, moderate content of sulphates and calcium, and high content of sodium (Table S2).

Fertilization solutions were prepared based on the substrate composition and the intake water analyses. The low nitrogen (LN) solution was prepared by adding 1.5 mM H3PO4 (Antonio Tarazona SL., Valencia, Spain), 4.85 mM K2SO4 (Antonio Tarazona SL., Valencia, Spain), 0.58 mM MgSO₄ (Antonio Tarazona SL., Valencia, Spain) plus 0.025 L/m3 of a microelements Welgro Hydroponic fertilizer (Química Massó S.A., Barcelona, Spain) containing boron (BO33-; 0.65% p/v), copper (Cu-EDTA; 0.17% p/v), iron (Fe-DTPA; 3.00% p/v), manganese (Mn-EDTA, 1.87% p/v), molybdenum (MoO42-; 0.15% p/v), and zinc (Zn-EDTA; 1.25% p/v). Normal nitrogen (NN) solution included the components listed above with the addition of 7.2 mM NH4NO3 to the intake water. The pH of the solutions was adjusted to 5.5-5.8 with 23% HCl (Julio Ortega SL., Valencia, Spain).

**Table S1.**
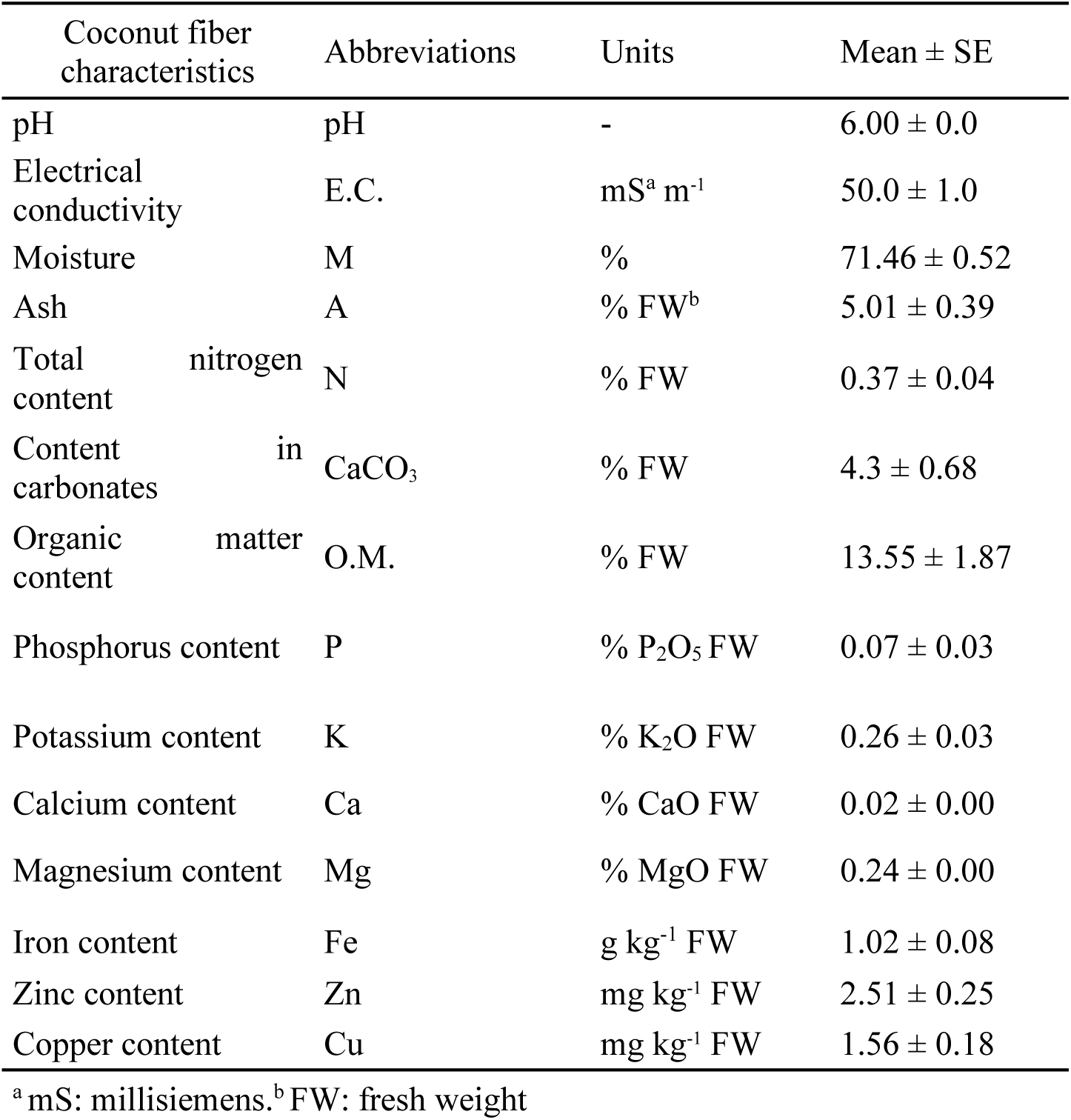
Mean values and standard error (SE) of coconut fiber substrate chemical composition used in the evaluation under low (LN) or normal (NN) of eggplant lines of *S. melongena* MEL5, MEL1 and MEL3 and its advanced backcrosses with *S. insanum*, *S. dasyphyllum* and *S. elaeagnifolium*.

**Table S2.** Chemical composition of intake irrigation water.

| Water characteristics | Abbreviations | Units | Intake water |
| --- | --- | --- | --- |
| pH | pH | - | 7.43 |
| Electrical conductivity | E.C. | mS cm<sup>-1</sup> | 1.4 |
| Nitrates | NO<sub>3</sub><sup>-</sup> | mmol/L | 0.29 |
| Nitrites | NO<sub>2</sub><sup>-</sup> | mmol/L | 0 |
| Ammonium | NH<sub>4</sub><sup>+</sup> | mmol/L | 0.03 |
| Phosphates | H<sub>2</sub>PO<sub>4</sub><sup>-</sup> | mmol/L | 0 |
| Sulphates | SO<sub>4</sub><sup>2-</sup> | mmol/L | 3.37 |
| Calcium | Ca<sup>2+</sup> | mmol/L | 3.79 |
| Magnesium | Mg<sup>2+</sup> | mmol/L | 0.38 |
| Potassium | K | mmol/L | 0.12 |
| Carbonates | CO<sub>3</sub><sup>2-</sup> | mmol/L | 0 |
| Bicarbonates | HCO<sub>3</sub><sup>-</sup> | mmol/L | 0.1 |
| Sodium | Na | mmol/L | 7.8 |
| Chlorine | Cl | mmol/L | 0.04 |

### 2.4. Phenotypic trait evaluation

Plants were evaluated for a total of 22 plant, fruit and composition traits (Table 1). A DUALEX® optical leaf clip meter (Force-A, Orsay, France) was used for measuring the chlorophyll, flavonol, anthocyanin contents and Nitrogen Balance Index (NBI®) in leaves^35, 36^. Data was obtained as the mean of 10 measurements in the upper and lower side of five leaves of each plant. At the end of the trial, stem diameter was measured with a caliper at the base of the stem and aerial biomass was immediately weighed after cutting the base of the stem with a Sauter FK-250 dynamometer (Sauter, Balingen, Germany). Subsequently, they were dried at room temperature, the leaves were separated from the stems, ground and weighed after drying in an oven at 70 °C to constant dry weight. The total number of fruits of each plant was harvested for determining yield. Nitrogen uptake efficiency (NUpE) was calculated as the total content of nitrogen (N) in fruit, stem and leaves divided by N supplied with the irrigation solution per plant; nitrogen utilization efficiency (NUtE) was calculated as total fruit yield in dry weight (yield [DM]) divided by the total content of N in fruit, stem and leaves and nitrogen use efficiency (NUE) was the result of the multiplication of NUpE and NUtE^6, 37, 38^.

**Table 1.** Plant, fruit, and composition traits evaluated in the *S. melongena* MEL1, MEL3 and MEL5 recurrent parents, and their respective advanced backcrosses individuals with *S. dasyphyllum*, *S. elaeagnifolium* and *S. insanum*, together with abbreviations and units used in the present study.

| Trait | Abbreviation | Units |
| --- | --- | --- |
| *Plant traits* |  |  |
| Chlorophyll leaf content | P-Chl | µg cm<sup>-2</sup> |
| Flavonol leaf content | P-Flav | µg cm<sup>-2</sup> |
| Anthocyanin leaf content | P-Anth | µg cm<sup>-2</sup> |
| Nitrogen Balanced Index | P-NBI | - |
| Aerial biomass | P-Biomass | kg FW<sup>a</sup> |
| Stem diameter | P-Diam | mm |
| Yield | Yield | g plant<sup>-1</sup> |
| Nitrogen Use Efficiency | NUE | - |
| Nitrogen Uptake Efficiency | NUpE | - |
| Nitrogen Utilization Efficiency | NUtE | - |
| *Fruit traits* |  |  |
| Fruit pedicel length | F-PedLength | mm |
| Fruit calyx length | F-CaLength | mm |
| Fruit length | F-Length | mm |
| Fruit width | F-Width | mm |
| Total number of fruits per plant | F-Number | - |
| Fruit mean weight | F-Weight | g |
| *Composition traits* |  |  |
| Nitrogen content in leaf | N-Leaf | g kg<sup>-1</sup> DM<sup>b</sup> |
| Carbon content in leaf | C-Leaf | g kg<sup>-1</sup> DM |
| Nitrogen content in fruit | N-Fruit | g kg<sup>-1</sup> DM |
| Carbon content in fruit | C-Fruit | g kg<sup>-1</sup> DM |
| Nitrogen content in stem | N-Stem | g kg<sup>-1</sup> DM |
| Carbon content in stem | C-Stem | g kg<sup>-1</sup> DM |

<sup>a</sup> FW: fresh weight. <sup>b</sup> DM:dry matter

For fruit traits, pedicel length, calyx length, fruit length and width were determined as the mean of at least three fruits per plant harvested at the commercially mature stage (i.e., physiologically immature). Fruits traits evaluated were measured with a caliper and their abbreviations and units are included in Table 1.

To determine fruit N and C content, at least five commercially mature fruits per plant were harvested, peeled, chopped and frozen in liquid N2 and stored at –80 °C. Subsequently, the frozen samples were lyophilized, ground until turned into fine powder and homogenized. Dry powder of leaves, stem and fruits was measured in samples of 0.5 g of freeze-dried powder. The analysis of N content was performed using the Dumas method with a TruSpec CN elemental analyzer (Leco, MI, USA). Carbon content was calculated from the measurements of carbon dioxide (CO2) using an infrared detector^39^. Certified reference standards of different N and C concentrations were used for the quantification.

### 2.5. Data analysis

For plant, fruit and composition data of each ABs set and the recurrent parents (*S. melongena* MEL5, MEL1 and MEL3) mean, standard deviation (SD), range values and coefficient of variation (CV, %) were calculated. Analysis of variance (ANOVA) was performed to detect significant mean differences between the two N cultivation conditions in the recurrent parents, and between each set of ABs and its corresponding recurrent parent in the LN conditions. Significant differences were detected with the Student-Newman-Keuls multiple range test at p < 0.05 using Statgraphics Centurion 18 software (StatPoint Technologies, Warrenton, VA, USA).

For each set of ABs and its recurrent parent cultivated in the same conditions (low nitrogen, LN) a principal component analysis (PCA) was performed. Pairwise Euclidean distances were calculated for the analysis of each PCA using R package stats^40^ of the R statistical software^41^. The PCA score and loading plots were drawn using R packages ggplot2^42^ and RColorConesa^43^. In addition, Pearson pair-wise correlation coefficients were calculated among traits for each set of ABs and *S. melongena* parents cultivated under LN conditions. Their statistical significance was evaluated using a Bonferroni correction at p < 0.01^44^ using R packages psych^45^ and corrplot^46^.

### 2.6. QTL detection and candidate gene identification

Detection of quantitative trait loci (QTLs) was performed for each set of ABs using the single QTL model for genome-wide scanning of the R package *R/qtl*^47^ of R statistical software v4.1.0^41^. The threshold of LOD score was established at the 0.05 probability level for significant QTLs. For each putative QTL detected, allelic effects were calculated by establishing significant differences between the means of each genotype with the Student-Newman-Keuls multiple range test (p < 0.05).

To identify potential candidate genes within each QTL region, a search was conducted using the ‘67/3’ eggplant reference genome assembly (V3 version)^14^. This search was performed through the Sol Genomics Network database (http://www.solgenomics.net).

## 3. Results

### 3.1. Genomic characterization

Genome coverage of the donor wild relatives in the whole sets of ABs was of 58.8% for *S. insanum*, 46.3% for *S. dasyphyllum*, and 99.2% *S. elaeagnifolium* when both heterozygous and homozygous introgressions are considered (Figure 1). Selection of ABs set of *S. insanum* prioritized a proper representation of chromosomes 1 (86.8%), 3 (80.9%), 6 (100%), 9 (78.4%), 10 (100%) and a region of chromosome 11 (34.8%) with several individuals per introgression for its evaluation (Figure 1A). The set of 59 selected ABs of *S. dasyphyllum* included most of chromosomes 1 (84.8%), 5 (86.3%), 6 (89.9%), 8 (89.1%) and. 12 (98.9%), plus an introgression at the end of chromosome 2 (11.5%) and another one at the beginning of chromosome 7 (20.2%) (Figure 1B). For the 59 ABs of *S. elaeagnifolium*, a high percentage of total genome coverage (in heterozygosis) was present for chromosomes 1, 2, 4, 6, 7, 8, 10 and 12 with several individuals for each large introgression (Figure 1C).

**Figure 1.**
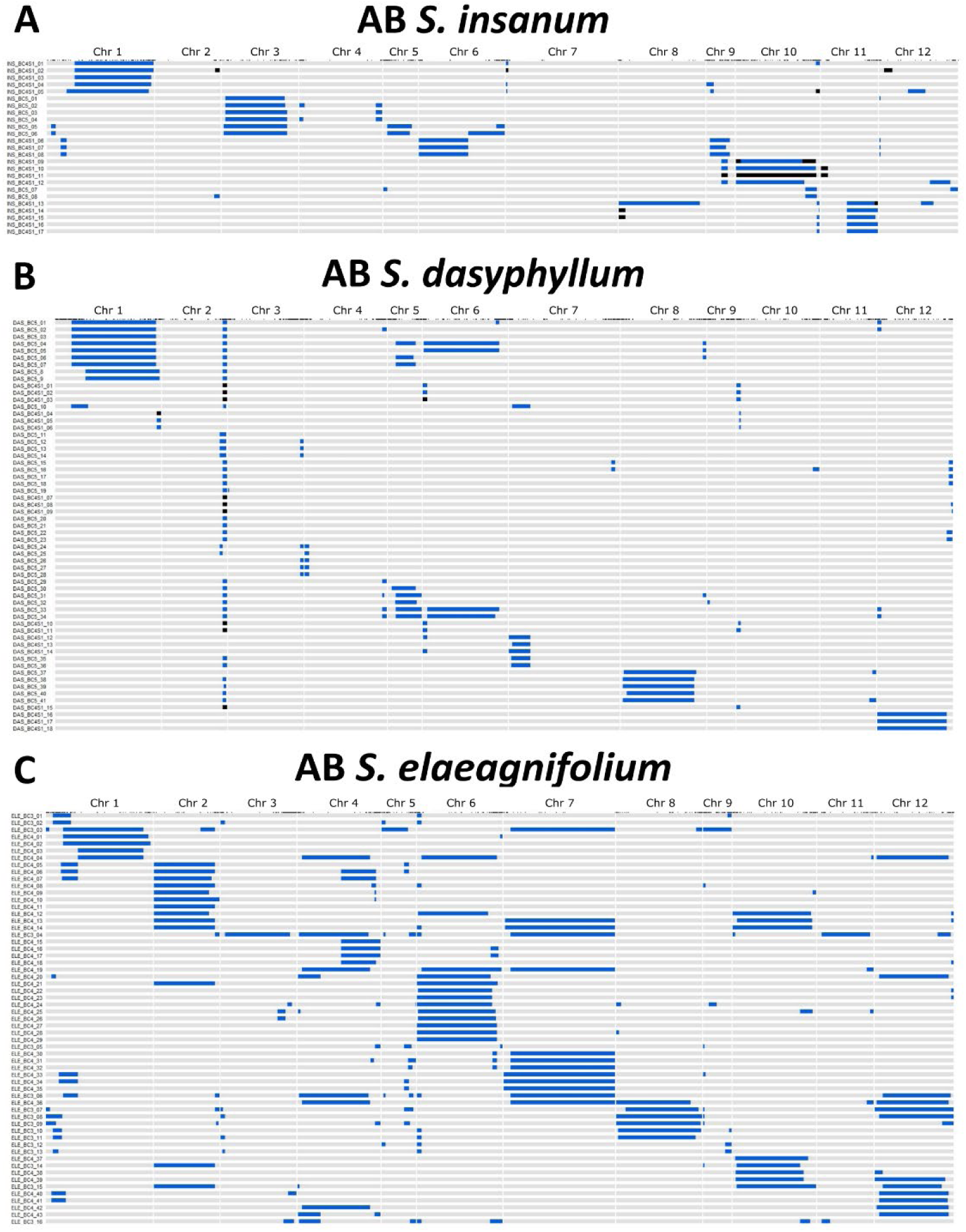
Graphical genotypes of advanced backcross (ABs) lines of *S. insanum* (A; n=25), *S. dasyphyllum* (B; n=59), and *S. elaeagnifolium* (C; n=59) assessed for the present experiment. Each row corresponds to the ABs codes and genotypes and the columns indicate the chromosomes. Heterozygous introgressions are colored in blue, homozyogous introgressions are colored in black and the genetic background of each recurrent parent (*S. melongena* MEL5, MEL1 and MEL3, respectively) are colored in grey.

The percentage of recovered genetic background from the recurrent parent is on average higher in the set of ABs of *S. insanum* (between 88.3% and 98.3%) and *S. dasyphyllum* (between 97.1% and 99.6%), while in the ABs of *S. elaeagnifolium* the percentage of recovery of the recurrent parent is lower (between 69.3% and 98.7%).

### 3.2. Characterization of recurrent parents and ABs

Overall, significant differences were detected between N treatments in *S. melongena* recurrent parents (MEL5, MEL1 and MEL3) for plant and composition traits, except for anthocyanin content in leaves (P-Anth) in MEL5 and carbon content in stem (C-Stem) in MEL1 (Tables 2, 3 and 4) *Solanum melongena* individuals cultivated under normal N conditions had higher chlorophyll content in leaf (P-Chl), nitrogen balanced index (P-NBI), aerial biomass (P-Biomass), stem diameter (P-Diam), yield (Yield), total number of fruits per plant (F-Number) and nitrogen and carbon content in leaves, fruits, and stem (N-Leaf, C-Leaf, N-Fruit, C-Fruit, N-Stem, C-Stem) than plants under low N conditions (LN). In addition, a significant decrease in values of flavonol and anthocyanin content in leaves (P-Flav, P-Anth), nitrogen use efficiency (NUE), nitrogen uptake efficiency (NUpE) and nitrogen utilization efficiency (NUtE) were observed in NN plants (Table 2). MEL5 showed the highest significant differences between treatments in yield (Yield, 3.7-fold) and total number of fruits (F-Number, 5.0-fold) with higher values under NN conditions, and the lowest differences in NUE (13.2-fold), NUpE (10.0-fold) and NUtE (1.3-fold) with highest values under LN conditions (Table 2). On the other hand, MEL1 presented the greatest differences between treatments for N content in the different plant parts, with higher values in NN than in LN for leaf (N-Leaf, 3.0-fold), fruit (N-Fruit, 2.0-fold) and stem (N-stem, 3.6-fold) (Table 4).

**Table 2.**
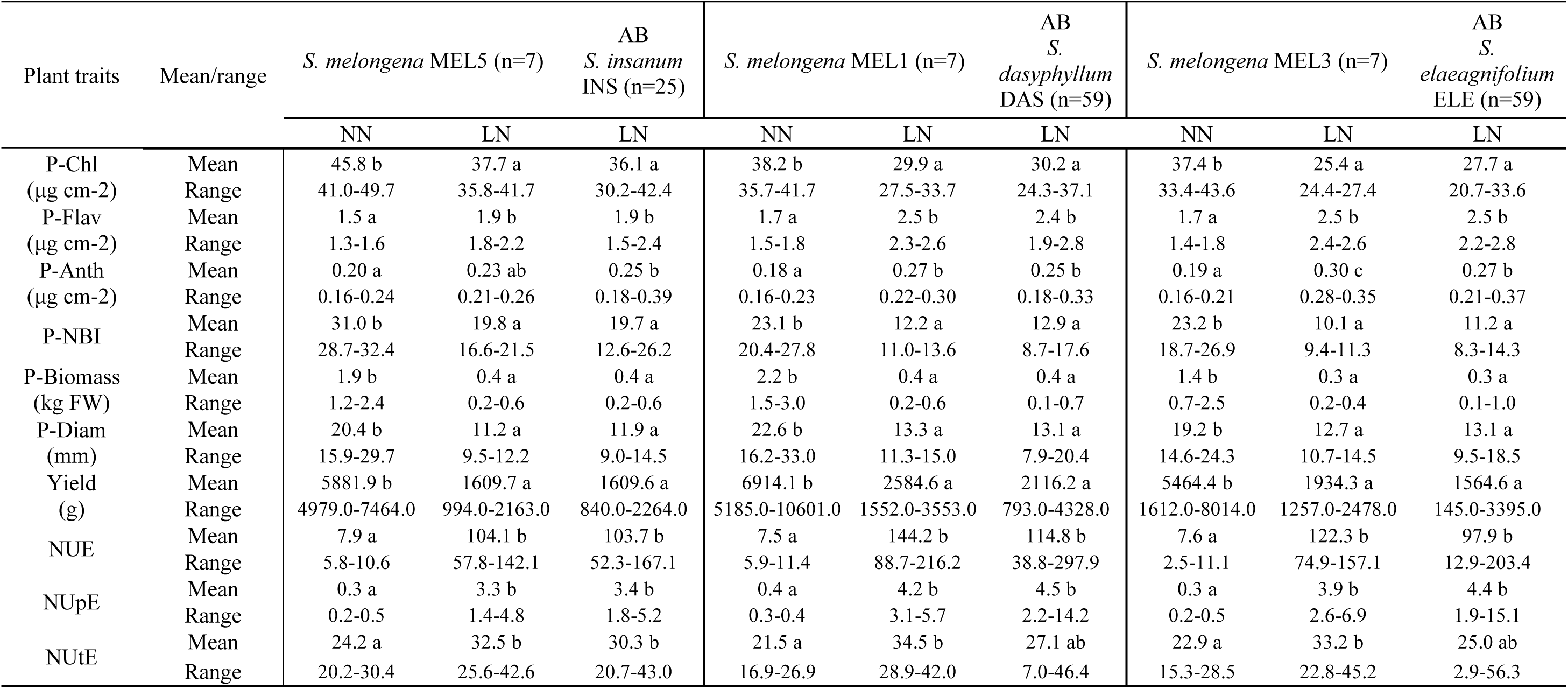
Mean values and range of plant traits of *S. melongena* MEL5, MEL1 and MEL3 in low nitrogen (LN) and normal nitrogen (NN) cultivation conditions and advanced backcrosses (AB) of *S. insanum* (INS; n=25), *S. dasyphyllum* (DAS; n=59) and *S. elaeagnifolium* (ELE; n=59) in low nitrogen (LN) cultivation conditions. The full name of each trait in the first column can be found in Table 1. For each trait, means with different letters are significant according to the Student–Newman–Keuls multiple range test (p < 0.05).

**Table 3.**
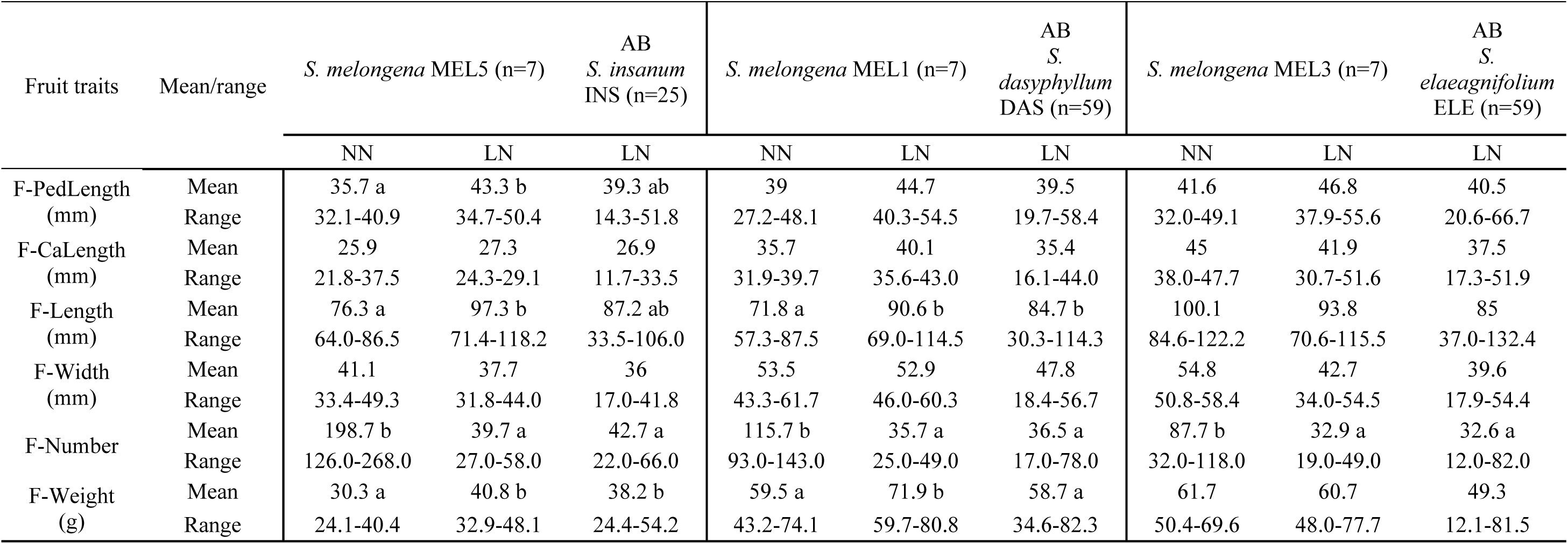
Mean values and range of fruit traits of *S. melongena* MEL5, MEL1 and MEL3 in low nitrogen (LN) and normal nitrogen (NN) cultivation conditions and advanced backcrosses (AB) of *S. insanum* (INS; n=25), *S. dasyphyllum* (DAS; n=59) and *S. elaeagnifolium* (ELE; n=59) in low nitrogen (LN) cultivation conditions. The full name of each trait in the first column can be found in Table 1. For each trait, means with different letters are significant according to the Student–Newman–Keuls multiple range test (p < 0.05).

**Table 4.**
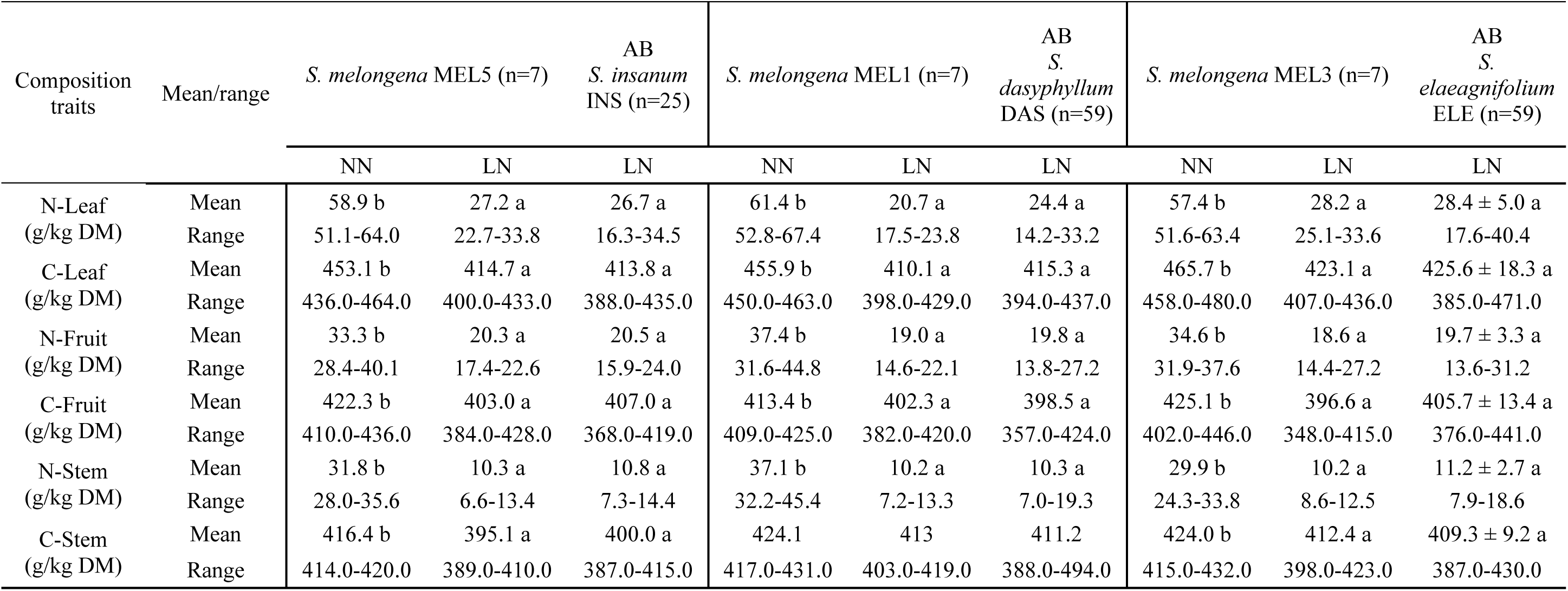
Mean values and range of composition traits of *S. melongena* MEL5, MEL1 and MEL3 in low nitrogen (LN) and normal nitrogen (NN) cultivation conditions and advanced backcrosses (AB) of *S. insanum* (INS; n=25), *S. dasyphyllum* (DAS; n=59) and *S. elaeagnifolium* (ELE; n=59) in low nitrogen (LN) cultivation conditions. The full name of each trait in the first column can be found in Table 1. For each trait, means with different letters are significant according to the Student–Newman–Keuls multiple range test (p < 0.05).

Some differences were observed between recurrent parents in N treatments for fruit shape and size traits (Table 3). Fruit pedicel length (F-PedLength) was statistically higher in MEL 5 cultivated under LN conditions than under NN conditions, and the same was observed for fruit length (F-Length) and fruit mean weight (F-Weight) in MEL1 and MEL5. For fruit calyx length (F-CaLength) and fruit width (F-Width) no statistically significant differences were detected (Table 3).

No significant differences were observed between each set of ABs and its corresponding recurrent parent cultivated under low N conditions for plant traits, except for anthocyanin content in leaf (P-Anth) in the set of ABs of *S. elaeagnifolium*, being the values significantly higher in MEL3 (1.1-fold) (Table 2). The same results were observed for fruit traits, except for mean fruit weight (F-Weight) in the set of ABs of *S. dasyphyllum*, which displayed significantly lower mean values than its recurrent parent MEL1 individuals in fruit mean weight (F-Weight; 1.2-fold) (Table 3). For composition traits, no statistically significant differences were detected (Table 4).

The distribution ranges for traits evaluated in the three sets of ABs were wider than those observed in the recurrent parents cultivated under low N conditions and transgressive individuals were found for all traits. The recurrent parents MEL5, MEL1 and MEL3 cultivated under NN conditions showed a wider distribution range for aerial plant biomass (P-Biomass), stem diameter (P-Diam) and yield (Table 2). Also, the same results were observed in MEL5 for F-Number and nitrogen content in fruit and stem (N-Fruit and N-Stem), in MEL1 for nitrogen content in stem (N-Stem), and in MEL3 for the total number of fruits (F-Number) (Table 3, Table 4).

### 3.3. Principal components analysis

A PCA was performed with the traits evaluated for each of the sets of ABs with *S. insanum*, *S. dasyphyllum* and *S. elaeagnifolium* (Figure 2). Three groups of traits can be observed in common, one includes chlorophyll content-related traits (P-Chl, P-NBI) and nitrogen content in plant (N-Leaf and N-Stem), another group of correlated traits includes plant vigor traits (P-Biomass, P-Diam), yield, nitrogen use efficiency (NUE) and the total number of fruits (F-Number), and the third group involves fruit size traits (F-PedLength, F-CaLength, F-Length, F-Width and F-Weight).

**Figure 2.**
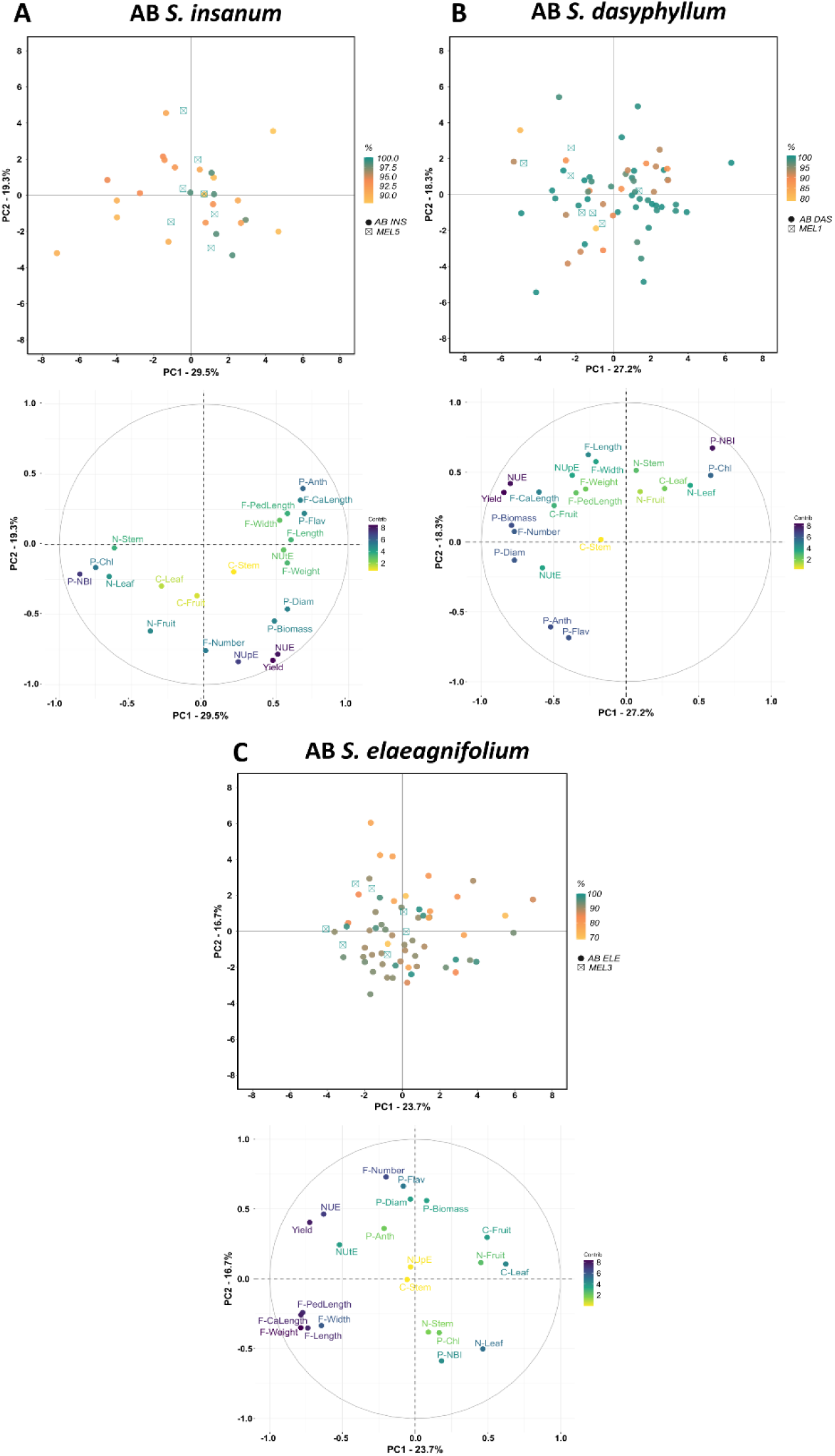
PCA score plot and loading plot based on the two first principal components of PCA performed for all traits of advanced backcrosses (ABs) of lines of *S. insanum* (A; n=25), *S. dasyphyllum* (B; n=59) and *S. elaeagnifolium* (C; n=59) and recurrent parents MEL5 (n=7), MEL1 (n=7) and MEL3 (n=7). The accessions are represented by different symbols according to the recurrent parent (MEL5, MEL1 and MEL3, respectively) and ABs of each line. Gradient of color in PCA score plot according to recovery percentage (%) from recurrent parent (100%: green to 80% (A); 70% (B); 90% (C): yellow). Gradient of color in PCA loading plot according to contribution proportion of each trait (8: dark blue to 2: yellow). The full name of each trait can be found in Table 1.

The PCA performed for the set of ABs of *S. insanum* and its recurrent parent *S. melongena* MEL5 revealed that the first two components accounted for 48.8% of the total variation observed, with PC1 and PC2 accounting for 29.5% and 19.3%, respectively (Figure 2A). The distribution of individuals in the PCA score plot showed that recurrent parent individuals of MEL5 were positioned along the central axis of PC1. Individuals with higher recovery percentages were located closer to the recurrent parentals. The first principal component displayed high negative correlation values with chlorophyll content-related traits (P-Chl, P-NBI), and positive correlations with P-Flav, P-Anth and fruit calyx length (F-CaLength). Nitrogen uptake efficiency (NUpE), yield, NUE and F-Number were highly negatively correlated with PC2, and P-Anth and F-CaLength were positively correlated to PC2 (Figure 2A, Table S3).

For the set of ABs of *S. dasyphyllum* and its recurrent parent *S. melongena* MEL1, the first and the second principal components (PCs) accounted for 27.2% and 18.3%, respectively, of the variation (Figure 2B). The projection of individuals in the PCA score plot showed that the individuals of the recurrent parent MEL1 displayed a wide distribution, being intermingled with some ABs individuals. Individuals with different recovery percentages were distributed all over the graph. The first component was highly negatively correlated with yield, NUE, plant vigor traits (P-Biomass, P-Diam) and F-Number, and positively with chlorophyll content-related traits (P-Chl, P-NBI). The second component was highly negatively correlated with flavonol and anthocyanin content in leaves (P-Flav and P-Anth), and positively with nitrogen balance index (P-NBI), and fruit length (F-Length) (Figure 2B, Table S3).

Regarding PCA performed for ABs of *S. elaeagnifolium* and its recurrent parent *S. melongena* MEL3, the first and the second components accounted, respectively, for 23.7% and 16.7%, of the observed variation (Figure 2C). The distribution of the individuals in the PCA score plot revealed a wide overall dispersion over the plot area, with most of the individuals with the lowest percentage of the recovered genetic background of the recurrent parent plotting apart from the recurrent parent MEL3 individuals. The composition traits carbon and nitrogen content in leaf (C-Leaf and N-Leaf) and carbon content in fruit (C-Fruit) were positively correlated with PC1, whereas some size-related fruit traits (F-Weight, F-CaLength, F-PedLength and F-Length) were highly negatively correlated with PC1. On the other hand, the second component was highly positively correlated with P-Flav and F-Number, and negatively correlated with P-NBI and N content in leaf (N-Leaf) (Figure 2C, Table S3).

**Table S3.**
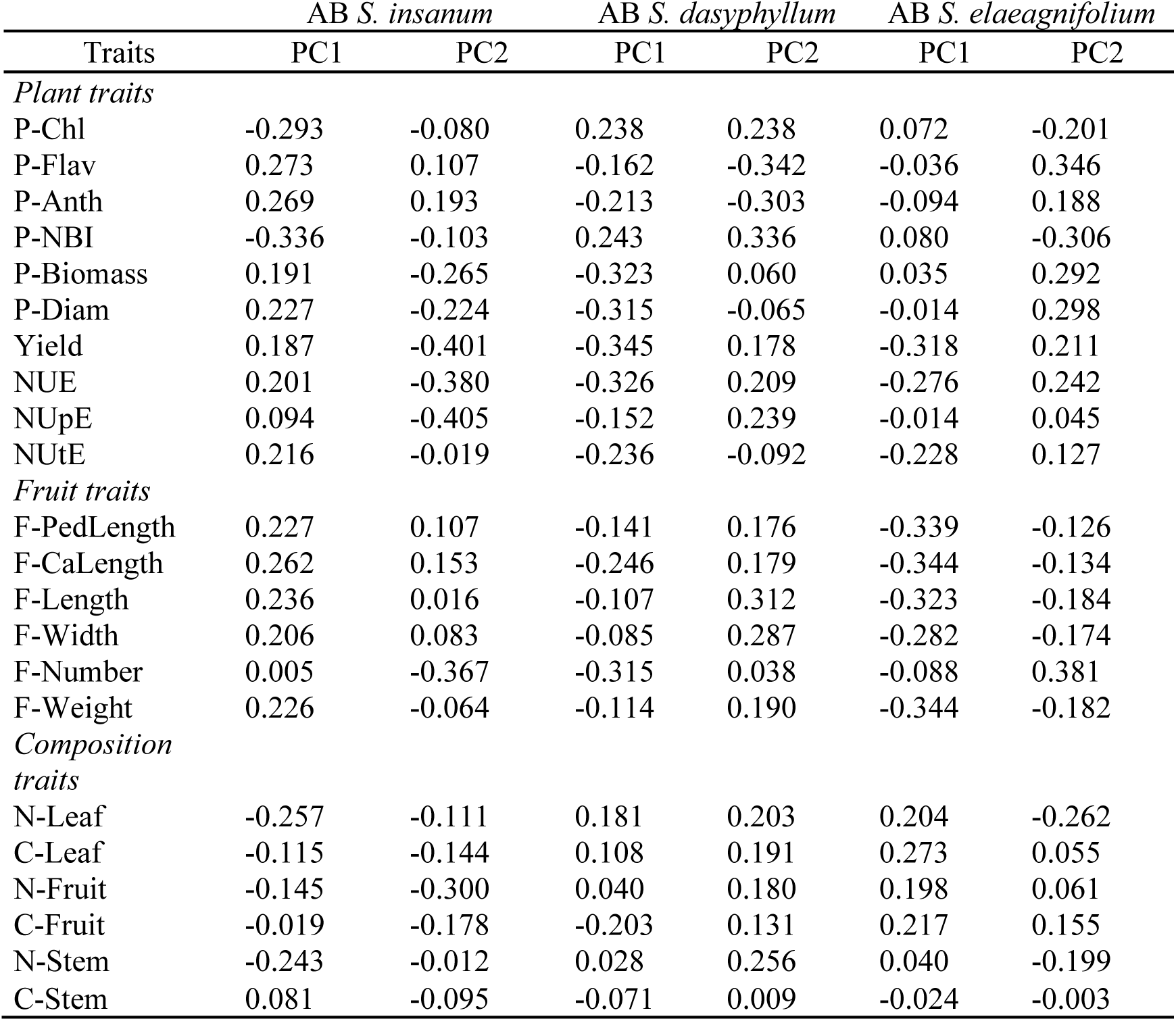
Correlation coefficients between traits evaluated and the two first principal components (PC1 and PC2) of the PCA of of advanced backcrosses (ABs) of lines of *S. insanum*, *S. dasyphyllum* and *S. elaeagnifolium* and its corresponding recurrent parents.

### 3.4. Correlations among traits

Significant Pearson’s linear correlations among traits evaluated were found in the three sets of ABs of eggplant with *S. insanum*, *S. dasyphyllum* and *S. elaeagnifolium* (Figure 3). For plant traits, negative correlations common to all ABs sets were observed among pigment content in leaves (P-Chl, P-Anth), and between nitrogen balance index (P-NBI) and anthocyanin and flavonol content in leaf (P-Anth, P-Flav). Positive correlations were detected between P-Chl and P-NBI. Traits related to plant vigor aerial plant biomass (P-Biomass) and stem diameter (P-Diam) were positively correlated (r > 0.6). In addition, yield, nitrogen use efficiency (NUE) and total number of fruits (F-Number) were positively correlated. Regarding fruit shape and size traits, shared positive correlations were found among fruit pedicel length (F-PedLength) and fruit width (F-Width) with fruit calyx length (F-CaLength) and fruit length (F-Length). For composition traits, leaf nitrogen content (N-Leaf) showed a common significant negative correlation with P-Flav (Figure 3).

**Figure 3.**
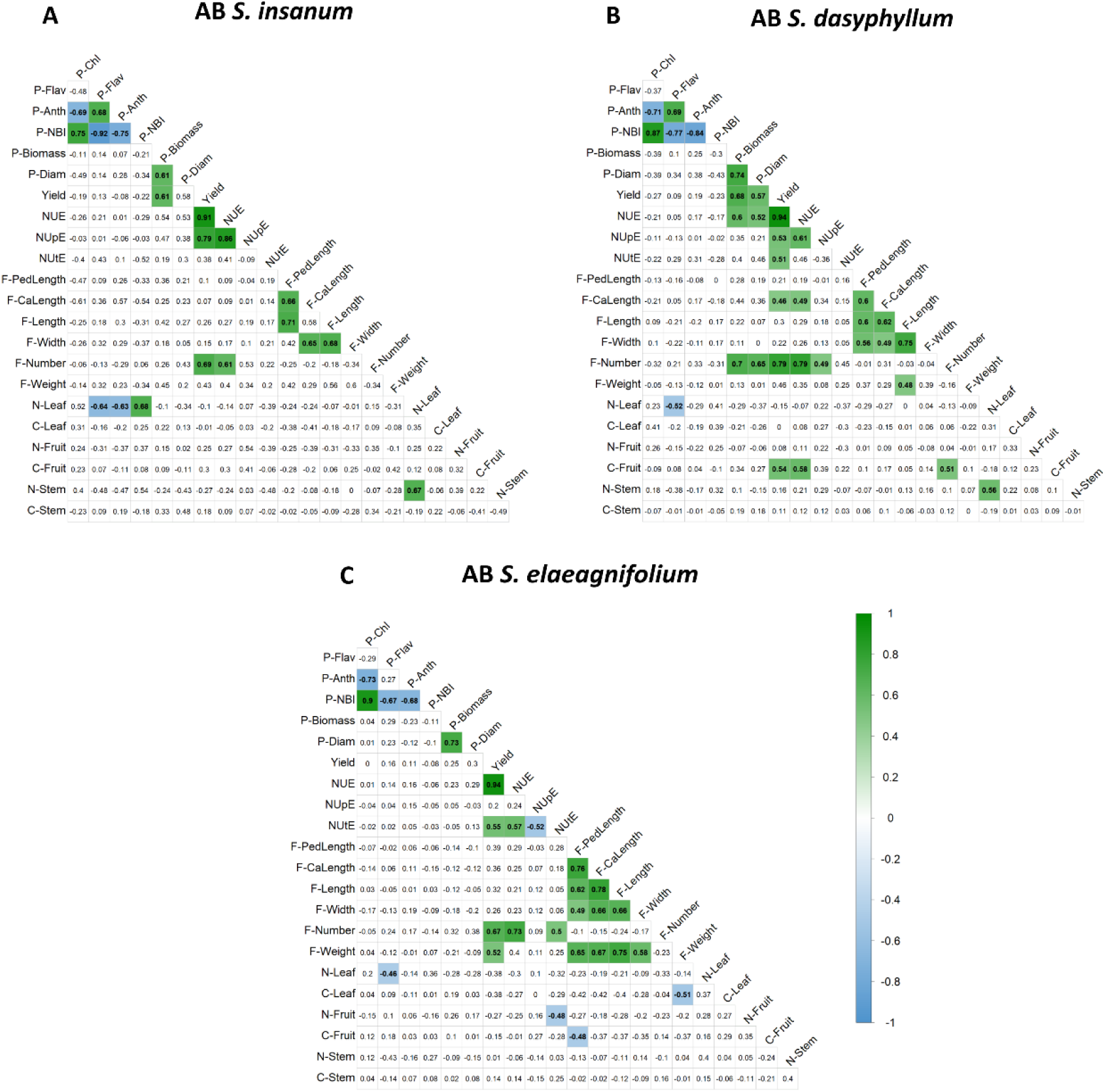
Pearson’s correlations among traits evaluated advanced backcrosses (ABs) of *S. insanum* (A; n=25), *S. dasyphyllum* (B; n=59) and *S. elaeagnifolium* (B; n=59). Only significant correlations at p<0.01 according to the Bonferroni tests are colored. Color scale from green (positive correlations) to blue (negative correlations). The full name of each trait can be found in Table 1.

The set of ABs of *S. insanum* shared different significant correlations with the set of ABs of *S. dasyphyllum*. In this way, a positive correlation was observed between P-Flav and P-Anth (r > 0.6) in both sets (Figure 3A, B). Yield, in addition to showing significant positive correlations with NUE, was also correlated with P-Biomass and nitrogen uptake efficiency (NUpE). Nitrogen content in leaves and stem (N-Leaf and N-Stem) also displayed a shared positive correlation in both ABs sets. On the other hand, several correlations among traits were shared between the ABs of *S. dasyphyllum* and the ABs of *S. elaeagnifolium*. In both sets, yield showed a positive correlation with nitrogen utilization efficiency (NUtE) (r > 0.5) (Figure 3B, C), while for fruit traits, F-length was positively correlated with F-CaLength and mean fruit weight (F-Weight), and F-PedLength was positively correlated with F-Width.

In addition, each set of ABs showed specific correlations. In this way, for ABs of *S. insanum*, N-Leaf displayed a positive correlation with P-Anth and a negative with P-NBI (Figure 3A). Regarding specific significant correlations found in ABs of *S. dasyphyllum*, NUE and F-Number were positively correlated with plant vigor-related traits (P-Biomass and P-Diam) (r > 0.7) (Figure 3B). NUE was also correlated with F-CaLength and carbon content in fruit (C-Fruit) and F-Number showed a positive correlation with NUpE. Additionally, yield showed positive correlations among F-CaLength and C-Fruit in this set of ABs. Finally, for ABs set of *S. elaeagnifolium*, NUtE showed positive correlations with other nitrogen use efficiency parameters (NUE and NUpE) and with F-Number, and a negative correlation with nitrogen content in fruit (N-Fruit) (Figure 3C). Mean fruit weight (F-Weight) was positively correlated with yield and traits related to fruit size (F-PedLength, F-CaLength, F-Length and F-Width), and negatively correlated with carbon content in leaf (C-Leaf). In addition, in this set of ABs F-PedLenght was negatively correlated with C-Fruit.

### 3.5. Detection and effect of putative QTLs

A total of 16 putative significant QTLs were found in the analysis of the three different sets of ABs (Table 5). Five QTLs (flavonol leaf content, *fl-9*; nitrogen balanced index, *nb-9*; fruit mean weight, *fw-9*; nitrogen content in leaf, *nl-9*; and nitrogen content in stem; *ns-9*) were found in ABs of *S. insanum* at the same position on chromosome 9 (Table 5, Figure 4A). QTLs for flavonol leaf content (*fl-9*) and for mean fruit weight (*fw-9*) presented similar effects (Figure 5A, C). In contrast, opposing effects were observed for QTLs associated with nitrogen balance index (*nb-9*), nitrogen leaf content (*nl-9*) and nitrogen stem content (*ns-9*), which presented significant higher values in individuals with heterozygous introgression (Figure 5B, D, E). Regarding ABs of *S. dasyphyllum*, eight of them were identified. Three QTLs were detected at the same position on chromosome 1 (Table 5, Figure 4B). QTL for flavonol leaf content (*fl-1*) presented significant higher values in individuals with heterozygous introgression, whereas an opposite QTL effect was observed for nitrogen balanced index (*nb-1*) (Figure 5G, H). For stem diameter (*di-1*), a dominance of the *S. melongena* allele decreasing the values was observed (Figure 5J). On chromosome 2, five QTLs were located at the same position (Table 5, Figure 4C). Chlorophyll leaf content (*ch-2*) presented dominance of the *S. dasyphyllum* allele, which displayed significant higher values while an opposite QTL effect was observed for biomass (*bi-2*) and yield (*yd-2*) (Figure 5F, I, K). For fruit pedicel length (*fp*-2) and fruit mean weight (*fd-2*) incomplete dominance was observed with a negative allelic effect of *S. dasyphyllum* introgression on the values (Figure 5L, M). The analysis of ABs of *S. elaeagnifolium* allowed the detection of two QTLs for fruit traits, namely fruit calyx length (*fc-2*) and fruit mean weight (*fw-2*) located at the same position on chromosome 2 (Table 5, Figure 4D), showing very similar effects with significant lower values in individuals with heterozygous introgression (Figure 5N, O). A QTL was also detected on chromosome 8 (Figure 4E), displaying significant higher carbon content in leaf (*cl-8*) corresponding to the *S. elaeagnifolium* heterozygous introgression (Figure 5P).

**Figure 4.**
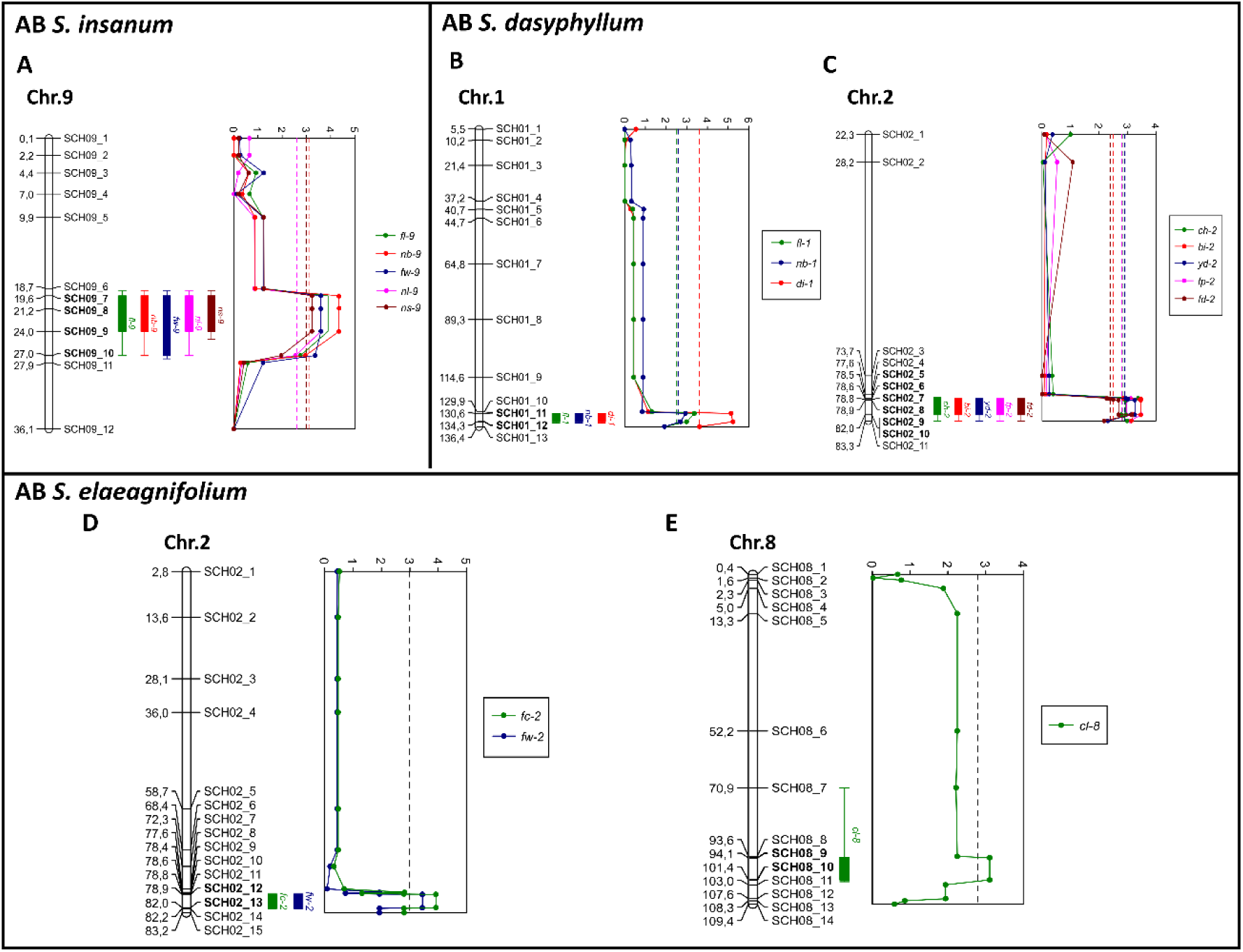
Physical map and LOD score of chromosomes with putative QTLs. A, ABs of *S. insanum*; B-C, ABs of *S. dasyphyllum*; and D-E, ABs of S*. elaeagnifolium*. Dotted lines indicate the LOD score thresholds of each QTL, with corresponding chromosome positions indicated on the physical map.

**Figure 5.**
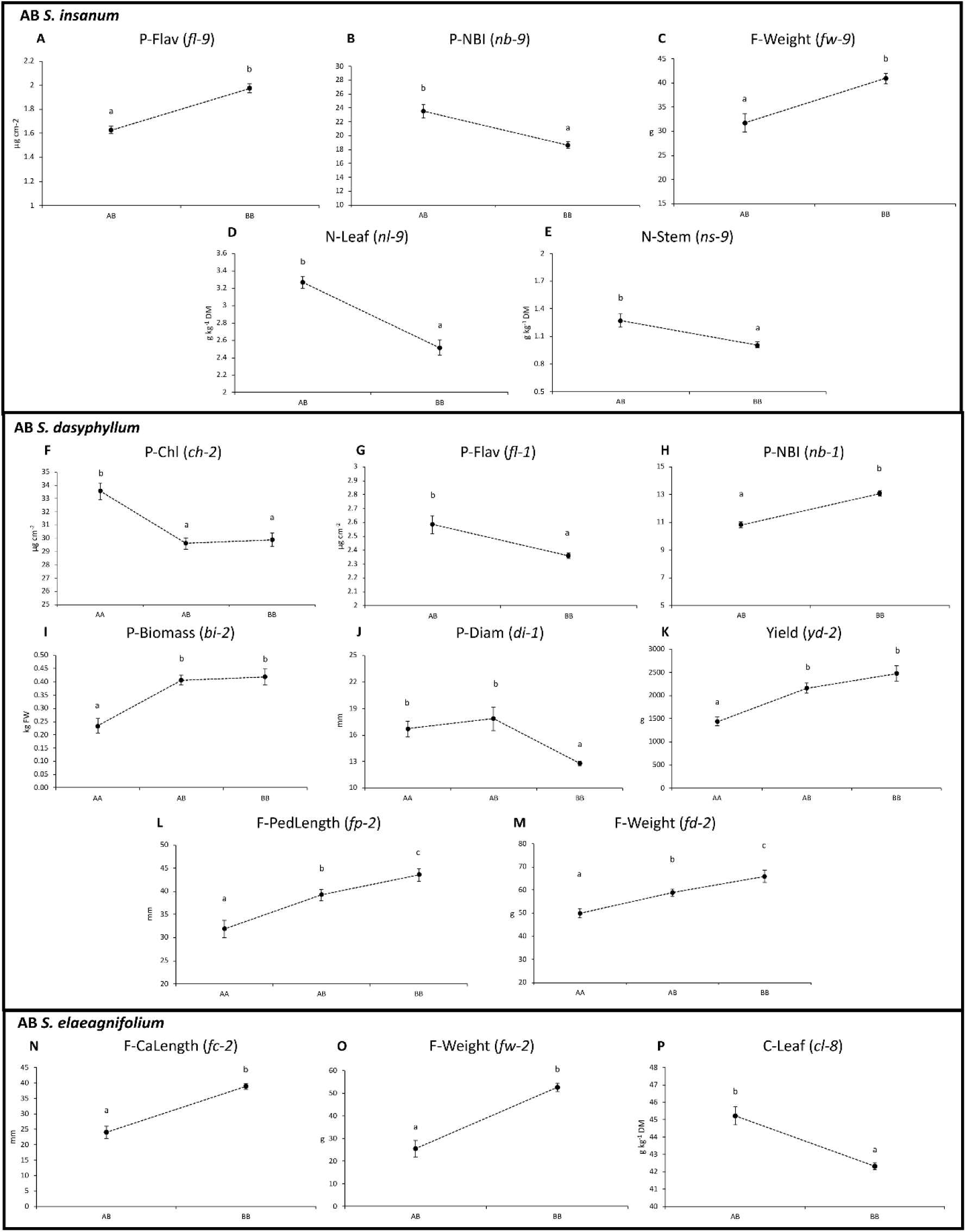
Effect plots for putative QTLs based on allelic distributions determined at the peak marker of the QTL. Three possible genotypes are indicated on the x-axis (A= wild parent, B= recurrent parent, AB= hybrid). A-E, ABs of *S. insanum*; F-M, ABs of *S. dasyphyllum*; and N-P, ABs of S*. elaeagnifolium*. The y-axis represents mean values of each trait for each genotype. Error bars represent standard error of the mean (SEM). For each trait, means with different letters are significant according to the Student–Newman–Keuls multiple range test (p < 0.05).

**Table 5.** List of putative QTLs detected for traits in advanced backcrosses (AB) of *S. melongena* (MEL5, MEL1 and MEL3) with *S. insanum*, *S. dasyphyllum*, *S. elaeagnifolium*. QTL name, chromosome, position (Mb) their genomic location, and LOD score.

| Advanced backcrosses (ABs) | Trait | QTL | Chr. | Position (Mb.) | LOD Score |
| --- | --- | --- | --- | --- | --- |
| *Plant traits* |  |  |  |  |  |
| ABs<br>*S. insanum* | Flavonol leaf content (P-Flav) | *fl-9* | 9 | 19.6-27.0 | 3.89 |
|  | Nitrogen Balanced Index (P-NBI) | *nb-9* | 9 | 19.6-27.0 | 4.35 |
|  | *Fruit traits* |  |  |  |  |
|  | Fruit mean weight (F-Weight) | *fw-9* | 9 | 19.6-27.0 | 3.6 |
|  | *Composition traits* |  |  |  |  |
|  | Nitrogen content in leaf (N-Leaf) | *nl-9* | 9 | 19.6-27.0 | 3.6 |
|  | Nitrogen content in stem (N-Stem) | *ns-9* | 9 | 19.6-27.0 | 3.23 |
| *Plant traits* |  |  |  |  |  |
| ABs<br>*S. dasyphyllum* | Chlorophyll leaf content (P-Chl) | *ch-2* | 2 | 78.5-83.3 | 3.38 |
|  | Flavonol leaf content (P-Flav) | *fl-1* | 1 | 130.6-134.3 | 3.37 |
|  | Nitrogen Balanced Index (P-NBI) | *nb-1* | 1 | 130.6-134.3 | 2.94 |
|  | Aerial biomass (P-Biomass) | *bi-2* | 2 | 78.5-83.3 | 3.47 |
|  | Stem diameter (P-Diam) | *di-1* | 1 | 130.6-134.3 | 5.23 |
|  | Yield | *yd-2* | 2 | 78.5-83.3 | 3.26 |
| *Fruit traits* |  |  |  |  |  |
|  | Fruit pedicel length (F-PedLength) | *fp-2* | 2 | 78.5-83.3 | 3.48 |
|  | Fruit mean weight (F-Weight) | *fd-2* | 2 | 78.5-83.3 | 3.15 |
| *Fruit traits* |  |  |  |  |  |
| ABs<br>*S. elaeagnifolium* | Fruit calyx length (F-CaLength) | *fc-2* | 2 | 78.4-83.2 | 3.92 |
|  | Fruit mean weight (F-Weight) | *fw-2* | 2 | 78.4-83.2 | 3.44 |
|  | *Composition traits* |  |  |  |  |
|  | Carbon content in leaf (C-Leaf) | *cl-8* | 8 | 94.0-101.4 | 3.11 |

### 3.6. Identification of candidate genes

The search for candidate genes in the ‘67/3’ eggplant reference genome assembly (V3 version)^14^ allowed identification of several potential candidate genes that may be associated to putative QTLs detected in this study. For the QTLs detected in ABs of *S. insanum* related to nitrogen content in plants, a gene encoding NITRATE TRANSPORTER 1/PEPTIDE TRANSPORTER (NRT1/PRT) FAMILY (NPF) proteins (SMEL_009g328470) was identified and mapped to the corresponding region of the detected QTL on chromosome 9. Regarding the QTLs detected on chromosome 2 associated with plant growth, yield, and fruit size parameters in ABs of *S. dasyphyllum* and *S. elaeagnifolium*, two distinct potential candidate genes were found to be possibly associated to these traits. One gene (SMEL_002g164700) encodes a PIN-FORMED (PIN) 8 auxin efflux transporter, while another gene (SMEL_002g167520) encodes a Myb-related protein 306 (MYB306), which is a transcription factor (TF) involved in anthocyanin regulation^48^. Additionally, within the same region on chromosome 2, a gene (SMEL_002g164340) was detected that encodes an NPF protein, which could also be associated with the same traits.

## 4. Discussion

The utilization of populations with CWR introgressions, such as advanced backcrosses, enables the utilization of variation present in the CWRs, facilitating the development of crop varieties that are suitable for sustainable agriculture practices^9^. Effective fertilizer management is crucial for improving sustainability, with nitrogen use efficiency (NUE) being a critical breeding goal due to its significant impact on economic and environmental factors^7, 49, 50^. Several studies have been conducted to improve NUE thought breeding in different crops, particularly in cereals and potato^51^. However, until recently, few efforts have been made to address this issue in eggplant recent years^31, 52–55^.

In this study, three sets of advanced backcrosses (ABs) of eggplant wild relatives from different genepools, namely *S. insanum* (GP1), *S. dasyphyllum* (GP2) and S*. elaeagnifolium* (GP3) were evaluated under the same conditions for the first time. The availability of eggplant ABs with introgressions in different chromosomes provided an overview of the potential of wild species for breeding when evaluated under low nitrogen input abiotic stress conditions. In addition, recurrent parental lines of *S. melongena* (MEL5, MEL1 and MEL3) were tested under low and normal N input, providing insights into the effect of N in the different traits in cultivated eggplant.

Previous studies evaluated nitrogen use efficiency (NUE) by conducting hydroponic culture in different Solanaceae species, including tomato^56, 57^, potato^58, 59^ and eggplant^52, 55^. Other works investigated NUE using soil cultivation methods^31, 60–62^, or combining different abiotic stresses^63–65^. Also, different approaches have been documented for the evaluation of NUE^66^. The use of pots and the automatic N fertilization system employed here allowed for more controlled conditions to evaluate the impact on different traits under low nitrogen input. This approach allows greater control of the experimental conditions, as all plants are subjected to the same fertilization and substrate, that soil cultivation.

The study revealed significant differences in plant and composition traits of *S. melongena* recurrent parents cultivated under different N treatments. Generally, plants grown under normal N conditions showed higher values in chlorophyll content and lower values in flavonol and anthocyanin content. These traits, which have been reported to correlate with nitrogen content in plant leaves, can be effectively measured using proximal optical sensors for nitrogen, thereby enabling optimized management of vegetable crop cultivation^67, 68^. The results also indicated that plants grown under normal N conditions displayed higher values for the studied traits such as aerial biomass, stem diameter, yield, total number of fruits per plant, and nitrogen and carbon content in plant and fruits. These findings are consistent with previous reports that demonstrated the impact of different nitrogen fertilization treatments on eggplant^53, 54, 69^. In contrast to Mauceri et al.^53^, the LN treatment resulted in much higher NUE, NupE and NUtE values than under NN. Similar results were found by Rosa-Martínez et al.^54^ suggesting that established fertilization practices may not always be the most efficient or sustainable approach to eggplant cultivation. Instead, carefully managed fertigation with reduced nitrogen inputs can improve NUE, resulting in higher yield per unit of N fertilization applied. Moreover, no significant differences were observed for most traits between each set of ABs and their respective recurrent parents under low N conditions. However, the wider distribution ranges for some traits observed within sets of ABs, which is in agreement with Villanueva et al.^31^, suggests that these materials may be of interest for enhancing the overall performance and variability of eggplant. The set of ABs used in this study do not cover the entire genome of wild eggplant relatives, indicating that further investigation into unexplored regions could result in significant findings for eggplant breeding under low nitrogen conditions.

The results of the PCA analysis revealed a relatively wide distribution of the advanced backcrosses in the PCA plot. Despite the general trend of AB individuals with lower recovery percentages being distributed separately from the recurrent parents, it is also observed that some accessions with high recovery percentages are distributed separately while others with low recovery percentages are closely located to the recurrent parents. This suggests that, in general, a high recovery of the recurrent genome background in advanced backcrosses is needed to have a general phenotype similar to the recurrent parent, although there are exceptions that may have great interest for breeding.

Breeding programs can benefit from correlations observed among similar traits in each set of ABs as these can help in predicting the phenotype of specific traits, requiring the assessment of fewer traits. Correlations established between traits evaluated by leaf clip meter Dualex® are consistent with results in previous studies^70, 71^. Similarly, associations identified between traits linked to plant vigor, such as plant biomass and stem diameter, and those related to yield, NUE and number of fruits per plant, as well as those associated with fruit morphology, are in agreement with several studies^31, 54, 72, 73^. Notably, some relevant correlations related to nitrogen content in plant were observed, including negative correlations between nitrogen content and flavonol and anthocyanin content in leaves, as well as positive intercorrelation between nitrogen content in leaves and stem. In addition, differences in correlations among NUE, NUpE and NUtE, and other traits, such as fruit calyx length or those related to plant vigor were, found between sets of ABs.

The identification of 16 putative significant QTLs across three different sets of ABs demonstrates the potential of genetic variation present in wild eggplant relatives. QTLs located on chromosome 2 for *S. dasyphyllum* and *S. elaeagnifolium*, associated with plant growth, yield and fruit size parameters, may suggest the presence of genetic linkage or a pleiotropic locus, which is consistent with results reported in previous studies^15, 54, 74, 75^. Furthermore, QTLs detected on chromosomes 2 and 9 for fruit pedicel and calyx length, as well as fruit weight, have also been identified in different collections and eggplant populations in earlier studies^15, 54, 74, 76–78^. For traits measured with the DUALEX® optical leaf clip meter, including chlorophyll, flavonol and NBI, novel QTLs in eggplant were identified on chromosomes 1 and 2 for *S. dasyphyllum*, and chromosome 9 for *S. insanum*. Additionally, for composition traits, a novel QTL for carbon content in leaf was found on chromosome 8 in ABs of *S. elaeagnifolium*, while two novel QTLs for nitrogen content in leaf and stem were located on chromosome 9 in ABs of *S. insanum*. In comparison, Rosa-Martínez^54^ reported QTLs for carbon leaf content on chromosomes 1, 5 and 10, and for leaf nitrogen content on chromosomes 4 and 9 in *S. melongena* introgression lines (ILs) with eggplant wild relative *S. incanum* as the donor parent. Interestingly, both studies detected QTLs associated with leaf nitrogen content on chromosome 9, suggesting a possible common underlying genetic factors.

In the identified QTL regions associated with plant growth, yield, fruit size and nitrogen-related parameters in ABs of *S. insanum*, *S. dasyphyllum*, and *S. elaeagnifolium*, several potential candidate genes have been detected. On chromosomes 2 and 9, genes encoding NITRATE TRANSPORTER 1/PEPTIDE TRANSPORTER (NRT1/PRT) FAMILY (NPF) proteins (SMEL_002g164340; SMEL_009g328470) were identified. These NPF proteins are involved in nitrate uptake and transport of various substrates in plants, contributing to diverse biological processes^79, 80^. Consequently, they may potentially influence nitrogen content and other nitrogen-related parameters in plants. Furthermore, on chromosome 2, a candidate gene (SMEL_002g164700) was identified, which encodes a PIN-FORMED (PIN) 8 auxin efflux transporter. This transporter is crucial for auxin distribution and affects a wide range of developmental processes in plants^81^. In the same chromosomal region, another candidate gene (SMEL_002g167520) was identified, encoding a Myb-related protein 306 (MYB306), This protein is a transcription factor (TF) involved in anthocyanin regulation^48^ and has been suggested to potentially influence both anthocyanin accumulation and fruit size in eggplant^82^.

## 5. Conclusions

This study highlights the potential of wild eggplant relatives for breeding under low nitrogen conditions by evaluating three sets of advanced backcrosses (ABs) and their recurrent parental lines. The findings reveal significant differences in plant, fruit, and composition traits in response to different nitrogen levels. Furthermore, we observed notable phenotypic variation among the ABs lines under low nitrogen fertilization, revealing the potential of introgression materials for genetic improvements in eggplant. The availability of genotyped lines with genetic variation allowed for the identification of putative QTLs. These insights may contribute to the development of breeding strategies aimed at improving eggplant productivity, quality, and nitrogen use efficiency under low nitrogen conditions, supporting sustainable agriculture practices.

## Acknowledgements

This work was supported by the project SOLNUE in the framework of the H2020 call SusCrop-ERA-Net (ID#47) and funded by Agencia Estatal de Investigación (PCI2019-103375) and by the Ministerio de Ciencia, Innovación y Universidades, Agencia Estatal de Investigación and Fondo Europeo de Desarrollo Regional (grant RTI2018-094592-B-I00 from MCIU/AEI/ FEDER, UE). The Spanish Ministerio de Ciencia e Innovación, Agencia Estatal de Investigación, and Fondo Social Europeo funded a predoctoral fellowship to Gloria Villanueva (PRE2019-103375). Pietro Gramazio is grateful to Spanish Ministerio de Ciencia e Innovación for a post-doctoral grant (RYC2021–031999-I) funded by (MCIN/AEI /10.13039/ 501100011033) and the European Union through NextGenerationEU/ PRTR.

## Data Availability Statement

Relevant data can be found within the paper and its supporting materials. All data of this study are available from the corresponding author upon reasonable request.

